# Visual properties of human retinal ganglion cells

**DOI:** 10.1101/766170

**Authors:** Katja Reinhard, Thomas A. Münch

**Author notes:** **Corresponding authors:** Katja Reinhard and Thomas A. Münch.

## Abstract

The retinal output is the sole source of visual information for the brain. Studies in non-primate mammals estimate that this information is carried by several dozens of retinal ganglion cell types, each informing the brain about different aspects of a visual scene. Even though morphological studies of primate retina suggest a similar diversity of ganglion cell types, research has focused on the function of only a few cell types. In human retina, recordings from individual cells are anecdotal or focus on a small subset of identified types. Here, we present the first systematic ex-vivo recording of light responses from 342 ganglion cells in human retinas obtained from donors. We find a great variety in the human retinal output in terms of preferences for positive or negative contrast, spatio-temporal frequency encoding, contrast sensitivity, and speed tuning. Some human ganglion cells showed similar response behavior as known cell types in other primates, while we also recorded light responses that have not been described previously. This first extensive description of the human retinal output should facilitate interpretation of primate data and comparison to other mammalian species, and it lays the basis for the use of ex-vivo human retina for in-vitro analysis of novel treatment approaches.

## Introduction

Vision starts in the retina, a highly structured part of the central nervous system. The retina performs important signal processing: the incoming images are captured by the photoreceptors, analyzed and split into parallel information streams by retinal circuits, and sent along the optic nerve to higher visual brain centers. Each of the parallel information streams is embodied by a type of ganglion cell and informs the brain about a particular aspect of the visual scene [1]. The non-primate mammalian retina contains over 40 of these different information streams, which can be distinguished based on both functional and morphological criteria [2–7].

One striking aspect of retinal architecture is that each ganglion cell type tiles the retina so that each feature can be extracted at each location in the visual field. Nevertheless, regional specializations do exist, for example the fovea of the primate retina, a region of very high visual acuity. The foveal region consists almost exclusively of four retinal ganglion cell types, the ON and OFF parasol cells, and the ON and OFF midget cells [8–10], which account for 50-70% of all ganglion cells in the primate retina [11]. Functional studies using human and non-human primates have often focused on these four most abundant retinal ganglion cell types [12–19]. Morphological studies of the complete primate retina, on the other hand, describe a similar variety in ganglion cell types as found in the non-primate retina with at least 17 morphologically identified types [11,20–23]. However, functional studies of these non-foveal ganglion cell types in non-human primates have been limited to a set of 7 types [14,24–29] and only midget and parasol cells have been recorded in human retina [18,19]. Additional physiological assessment of the human retina on the level of individual cells is anecdotal [30,31].

In this study, we present a survey of ganglion cell function in the non-foveal human retina. We performed multi-electrode array (MEA) recordings on *ex-vivo* retinas obtained from enucleation patients and recorded light-driven activity from dozens of human ganglion cells in parallel. MEAs have been successfully used in previous studies to characterize the retinal output in various animal models [4,12,32–37]. Our data represents the first systematic and non-selective recording and characterization of light responses from a large population of ganglion cells in human retina. In addition to providing an overview of the spectrum of light responses in the human retina, we compare the representation of the spatio-temporal stimulus space by human ganglion cells with published data from non-human primate retina and results from psychophysical studies.

## Results

### Recording light responses from donated human retinas

To record light responses from human retinal ganglion cells, we obtained human retinas from patients who had to undergo enucleation of one eye due to a uveal tumor. Retinal pieces (~ 3 × 3 mm^2^) were placed ganglion cell-side down onto multi-electrode arrays and responses to a set of light stimuli were recorded at photopic light intensities. Individual stimuli (gray-scale images) spanned at most 3 log units of brightness. Spikes were assigned to individual units (presumably retinal ganglion cells) during an offline, semi-manual spike sorting process based on principal component analysis of spike waveforms. Only clearly sortable units were considered for analysis (see Method section for details). In total, we obtained the spiking activity of 342 light-responsive single units in 15 retinal pieces obtained from 10 human retinas (Table 1).

**Table 1:** Human retinas used for this study. 15 ex-vivo human retinas were obtained. The table contains information about the donor (sex, age, known medical history), surgery conditions (the ischemia duration, i.e. time without oxygen and nutrient supply, and whether a dark lens was put on the donor’s eye during surgery), and the retina (left/right eye, part of the retina without tumor). During preparation, the retina would sometimes roll up immediately after vitrectomy (rolling) or the vitreous was sticking strongly to the retina (sticky). The last two columns indicate how many retinal pieces were used per retina for the final analysis and whether any light responses were detected in our recordings. Light gray rows: retinas with few light responses, not used for analysis. Dark gray rows: no detectable light responses. Bold: potential reasons for low quality. Ventr./v = ventral, dors./d = dorsal, temp./t = temporal, n = nasal, m = male, f = female, radiation = radiation of the tumor-bearing eye, detachment = partial retinal detachment prior to surgery, resp. = responses. *retina prepared by another group during a different study.

| Donor |  |  |  | Surgery conditions |  | Retina |  | Experiment notes |  |  |
| --- | --- | --- | --- | --- | --- | --- | --- | --- | --- | --- |
| ID | Sex | Age | Notes | Ischemia (min) | Dark lens | Eye | Retinal part (ventr./dors./temp./nasal) | Preparation | # Analyzed pieces | Any light resp.? |
| 1 | m | 72 | diabetic | 7 | - | right | vt | easy | 1 | yes |
| 2 | f | 49 | **radiation** | 7 | - | right | dn |  | - | - |
| 3 | f | 72 | diabetic | 7 | - | left | dn | rolling | 1 | yes |
| 4 | f | 69 | detachment | 17 | - | left | n | easy | 1 | yes |
| 5 | m | 53 |  | 7 | - | left | t | sticky | 1 | yes |
| *6 | m | 75 | sinus tumor | 18 | - |  |  |  | - | - |
| 7 | m | 89 |  | 25 | - | right | dn | sticky | - | yes |
| 8 | f | 42 |  | 20 | - | left | vt | sticky | - | - |
| 9 | f | 83 |  | 10 | - | right | t | sticky | 1 | yes |
| 10 | f | 49 |  | 10 | - | left | t | sticky | 3 | yes |
| 11 | f | 60 |  | 7 | yes | left | vn | sticky | 3 | yes |
| 12 | f | 74 | macular edema | 7 | yes | left | dt | easy | 2 | yes |
| 13 | m | 74 |  | 7 | yes | left | v | **very sticky** | - | yes |
| 14 | f | 79 | radiation 10y ago | 8 | - | left | n | easy | 1 | yes |
| 15 | m | 67 | detachment | 10 | - | left | n | easy | 1 | yes |

### Response properties across the population of ganglion cells

We aimed at characterizing the diversity of the output of the human retina with different visual stimuli (Fig. 1). We used drifting-grating stimuli to characterize the encoding of the spatio-temporal space (Fig. 1A). Of the 342 light-responsive cells, 86% responded to these stimuli. As a population, the recorded cells responded to a large spatio-temporal stimulus space including all tested spatial frequencies (100-4000 μm spatial period on the retina, corresponding to 2.66-0.07 cycles per degree (cyc/°)) and temporal frequencies (1-8 Hz) with an overall preference for stimuli of 500-4000 μm retinal size (0.53-0.07 cyc/°) and moving with 2-8 Hz (Fig. 1E). Figure 1E shows the response strength averaged across all recorded cells to the 24 different sinusoidal drifting gratings. To obtain the displayed heat-map, the amplitude in the Fourier Transform of the cells’ responses at the stimulus frequency was taken as response strength and normalized for each cell across the 24 grating stimuli. The distribution of preferred spatial and temporal frequencies per cell are shown in Figure 1F (maximum out of the 24 drifting-grating combinations). While the recorded ganglion cells showed responses to a broad range of spatial and temporal frequencies (Fig. 1E), they mostly responded best to coarse gratings (Fig. 1F left) and higher temporal frequencies (Fig. 1F right).

**Figure 1:**
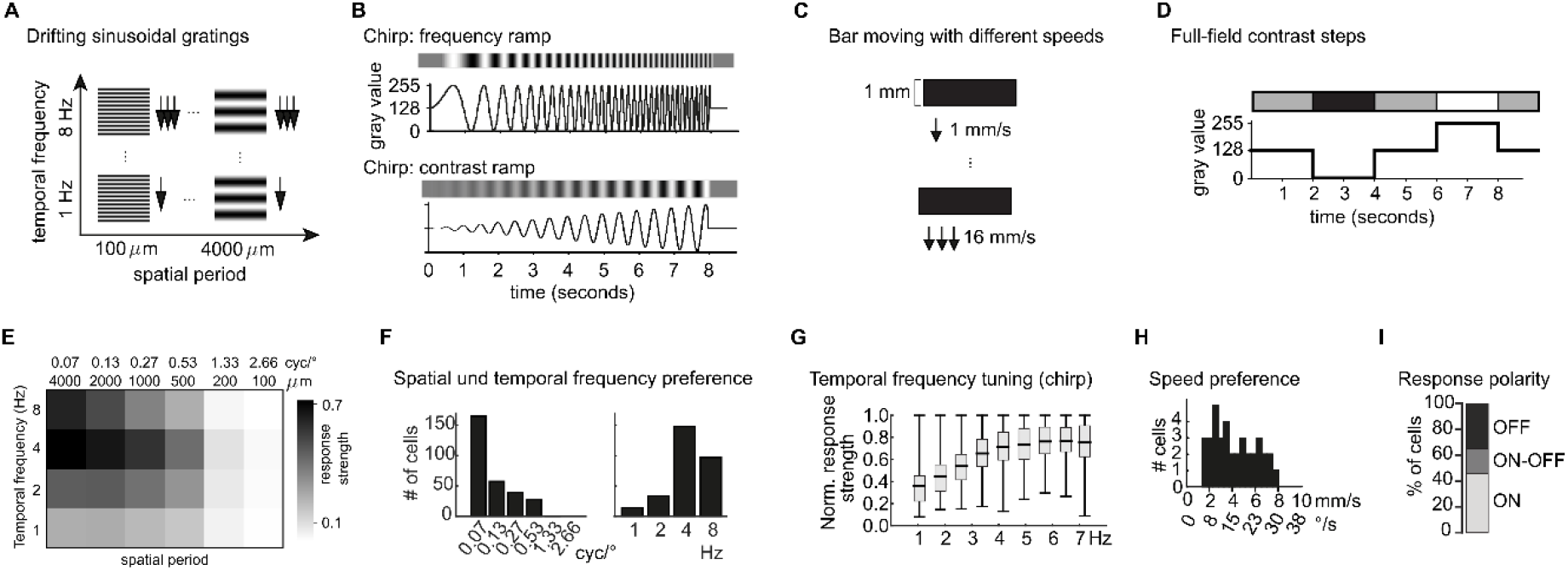
Population response of human retinal ganglion cells. Four different types of stimuli were used (A-D). A) Drifting gratings with 4 temporal frequencies and 6 spatial frequencies. B) Full-field “chirp” frequency ramp from 0.5 to 8 Hz and full-field “chirp” contrast ramp. C) Bar moving with 6 different speeds. D) Full-field flash contrast steps consisting of two positive and two negative contrast steps. (E) Response strength (amplitude of the Fourier Transform (FT) normalized to maximal response) to 24 drifting sinusoidal gratings with different spatial and temporal frequencies, averaged across N = 293 cells. (F) Distribution of preferred spatial (left) and temporal (right) frequencies in response to drifting gratings, N = 293 cells. (G) Normalized response strength (FT_response_/FT_stimulus_) to a full-field frequency ramp (“chirp”), in different frequency bands (Box-whisker-plots: mean, quartiles, maximum and minimum; N = 141 cells). (H) Distribution of the median preferred speed measured with a single moving bar, N = 37 cells. (I) Proportion of ganglion cells responding to positive full-field contrast steps (ON), negative contrast steps (OFF) or both (ON-OFF), N = 121 cells.

Temporal frequency preferences were further measured with a full-field frequency ramp (“chirp” stimulus, Fig. 1B top) which has proven to be an excellent stimulus to classify the behavior of retinal ganglion cells [2], and which drove activity in 41% of our analyzed cells. Here, response strength was defined as the ratio of the Fourier Transform of the cells’ response and the Fourier Transform of the stimulus. We analyzed this normalized response strength across all cells with chirp responses in discrete 1 Hz bins. Figure 1G shows the distribution of the normalized response strength for each bin across all responding cells (mean, quartiles and extremes). While there is at least one cell with a maximum response for each frequency bin, this chirp stimulus confirmed a general preference of the human retinal output for higher temporal frequencies. Bars moving with different velocity (Fig. 1C) were used to test for the preferred speed of ganglion cells and elicited clear responses in 11% of all cells. The distribution of the median preferred speeds (50% of the cumulative sum of the response amplitudes) was rather wide, ranging from bars moving between 2 and 8 mm/s (7.5 to 30 °/s) in different ganglion cells (Fig. 1H). Finally, response polarity was tested with full-field contrast steps (Fig. 1D). Over a third of the recorded cells responded consistently to this stimulus. Of those cells, 46% responded solely to positive full-field contrast-steps (ON-responses), 35% showed responses to negative contrast steps (OFF-responses), and the remaining 19% responded to both (ON-OFF-responses; Fig. 1I).

### Diversity in the output of human retina

One hallmark of the retina is the separation of visual information into different information streams embodied by distinct ganglion cell types. When analyzing individual cells, we found a wide range of response properties to our set of light stimuli, illustrated with 15 example cells in Figure 2. These example cells span the range of observed response polarity and transiency, spatio-temporal preferences, contrast sensitivity, and responsivity to local stimuli. For the following description, we group these cells based on their responses to full-field contrast steps (column 1 in Fig. 2), with cells responding only to positive contrast steps (ON-cells) in Figure 2A, cells responding exclusively to negative contrast steps (OFF-cells) in Figure 2B, and cells responding to both (ON-OFF cells) in Figure 2C.

**Figure 2:**
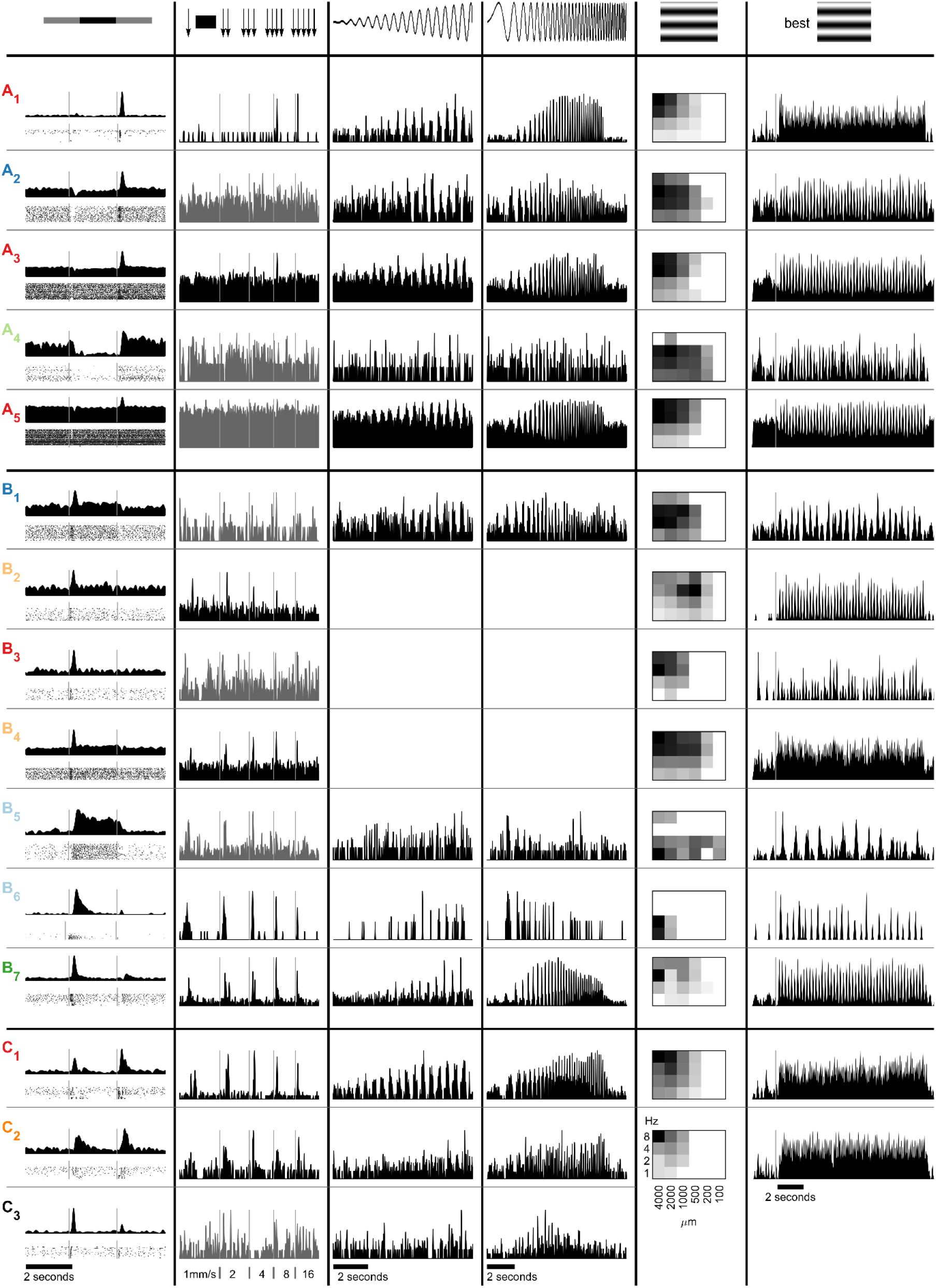
Example response properties of human ganglion cells. A) ON-cells, B) OFF-cells, C) ON-OFF-cells. Column 1: average firing rates (top) and raster plots (bottom) to full-field contrast steps; column 2: response to bar moving with different speeds; column 3: activity during a full-field contrast ramp at 2 Hz; column 4: response to “chirp stimulus” (full-field temporal frequency modulation from 0.5 to 8 Hz); column 5: normalized response strengths to 24 sinusoidal drifting-gratings; column 6: firing rate for sinusoidal drifting-grating with maximal response (white square in column 5). Stimuli are depicted on top. Colors correspond to Figure 4. White space indicates that the stimulus was not presented to this cell. Firing rates plotted in gray indicate that the cell did not respond consistently to this stimulus.

#### ON-cells

The cell A_1_ in Figure 2 showed a very transient response to a positive contrast step (at the transition from black to gray full-field stimulation, column 1). It preferred coarse drifting-gratings with a high temporal frequency (columns 5 and 6), which is consistent with the steep increase in responsivity when shown a spatially homogeneous frequency ramp (column 4). The cell also responded to fast local stimulation by a moving bar (column 2). When probed with a contrast ramp, the activity of this cell was already modulated at relatively low contrast (column 3). A_2_ and A_3_ are additional transient ON-cells.

Compared to the first cell, these cells responded well to a broader spectrum of temporal frequencies (columns 4 and 5), but cell A_2_ did not show significant activity modulation in response to a moving bar (column 2).

Some recorded ganglion cells had very high spontaneous firing rates such as the cell A_4_ and A_5_. Nevertheless, they precisely encoded various combinations of temporal and spatial frequency stimuli and showed selective activity modulations to their preferred contrast step. In addition, some cells like example cell A_4_ were strongly inhibited by negative contrast.

#### OFF-cells

Four examples of transient OFF-cells are shown in Figure 2B_1-4_. Only cell B_2_ and B_4_ responded consistently to moving bars, but with opposite preferences (B_2_ prefers slow bars, B_4_ fast bars). Their speed preferences are also reflected in their responses to drifting gratings. Both cells prefer similar temporal frequencies (4-8 Hz), but different spatial frequencies resulting in distinct speed preferences (1-2 mm/s for B_2_ and 16-32 mm/s for B_4_). B_3_ responds only to fast and wide stimuli and not to moving bars, while B_1_ responds to a broad range of spatial frequencies and slower stimuli.

The sustained cells B_5_ and B_6_ both preferred low temporal frequencies (1-2 Hz) when probed with a drifting-grating stimulus or the chirp stimulus. The less sustained cell B_6_ responded strongly to all moving bars, while the cell B_5_ did not respond consistently to this stimulus. Finally, the OFF-cell B_7_ exhibited a rebound or delayed response to positive contrast after an initial inhibition. The cell showed a preference for temporal frequencies around 4 Hz and higher contrast stimuli, and it responded well to all speeds of a moving bar.

#### ON-OFF-cells

ON-OFF cells may show rather sustained (cells C_1_ and C_2_) or transient responses (C_3_). Some clearly prefer higher temporal frequencies (C_1_), others responded only to low frequencies (C_3_). While the cell C_1_ responded well to the whole contrast ramp, the other two example cells showed some activity modulation only to maximal contrast. Interestingly, the cell C_3_ responded well to temporal frequencies around 3 Hz when exposed to full-field stimulation (chirp stimulus) but did not respond to moving bars.

### Spatio-temporal properties of human ganglion cells correspond to psychophysical detection threshold

The responses to drifting or sign-inverting grating stimuli have often been used to characterize, identify, and compare different retinal ganglion cell types. We therefore explored the spatio-temporal stimulus space encoded by the human retina in more detail. The heat-map in Figure 1E (replicated in Fig. 3A) indicates that the mid-peripheral human retina responds well to all presented temporal frequencies and shows a general preference for coarser stimuli. To directly compare the human retina responses to published psychophysics and non-human primate data, we computed the spatial response curve of the whole population of recorded cells (Fig. 3A top and 3B). This was achieved by normalizing every cell’s responses to each spatial frequency presented at its optimal temporal frequency, and then averaging these individual spatial response curves. In the corresponding way, the average temporal response curve was calculated (Fig. 3A left and 3C). The average spatial response curve dropped below 10% of its maximum for stimuli of 1.55 cyc/° and finer (Fig. 3A top). This *in-vitro* spatial threshold corresponds well to previously determined psychophysical detection thresholds in the mid-peripheral visual field (4 cyc/° at 14° visual angle and 2 cyc/° at 30°) [38].

**Figure 3:**
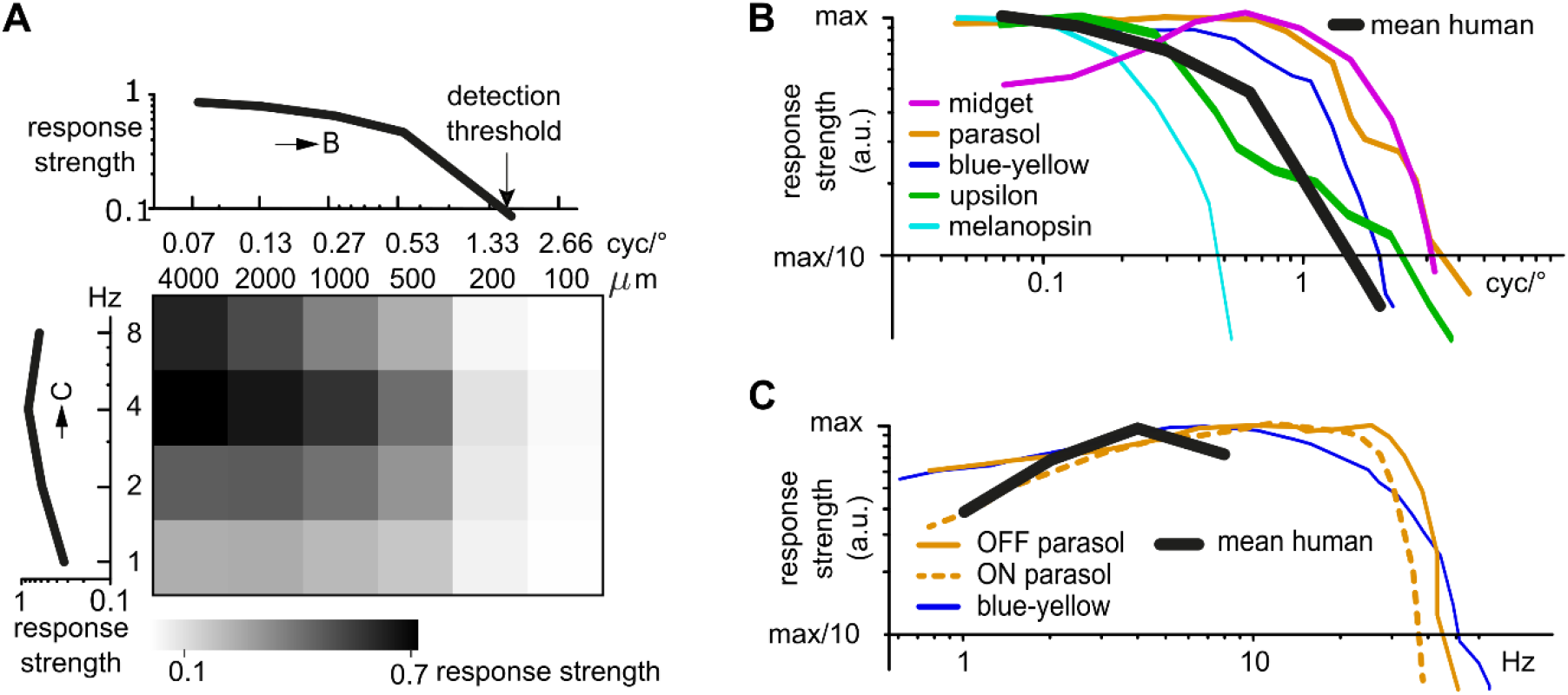
Human retinal ganglion cells show similar spatial and temporal frequency response curves as non-human primate retinal ganglion cells. *(A) Heat map: Average responsivity of human retinal ganglion cells for drifting sinusoidal gratings, replicated from Fig. 1E (N* = *293). Curves: Spatial frequency (top) and temporal frequency (left) response curves (mean across all cells). (B, C) Spatial and temporal response curve in comparison with published data on non-human primate ganglion cells. Non-human primate data adapted from* [39] *(midget);* [40,41] *(parasol);* [40,42] *(blue-yellow);* [24] *(upsilon);* [27] *(melanopsin).*

### Comparison to non-human primate data

Temporal and spatial frequency preferences have been used as the main parameter in several studies on non-human primate retina to characterize and identify different ganglion cell types [24,27,39–42]. In all non-human primate publications considered here for comparison with our human ganglion cell data, response strength has been given either as absolute number of spikes or as a normalized amplitude of the Fourier Transform of the cells’ responses. We extracted the response curves from these publications (midget ganglion cells [39]; parasol ganglion cells [40], [43]; blue-yellow ganglion cells [40,42]; upsilon ganglion cells [24]; melanopsin ganglion cells [27]) and overlaid them with the population tuning curves obtained from our human ganglion cell data, as shown in Figure 3B and 3C. Both the average spatial response curve (Fig. 3B) and the average temporal response curve (Fig. 3C) for the human retina lie within the range of published data from different primate ganglion cell types.

### Spatio-temporal clusters of human ganglion cell responses

No stimulus induced responses in more cells than the drifting-grating stimulus (n = 293 responding cells). In addition, the response properties of the example cells in Figure 2 suggest a great variety in the spatio-temporal preferences of the recorded human ganglion cells. Proper classification of the recorded cells would require morphological information and/or denser electrophysiological recordings to reveal mosaic information. However, to get a more systematic handle on the response diversity, we used k-means clustering to cluster the responses to drifting-grating stimuli of the recorded ganglion cells into 10 groups of cell types (Fig. 4A).

**Figure 4:**
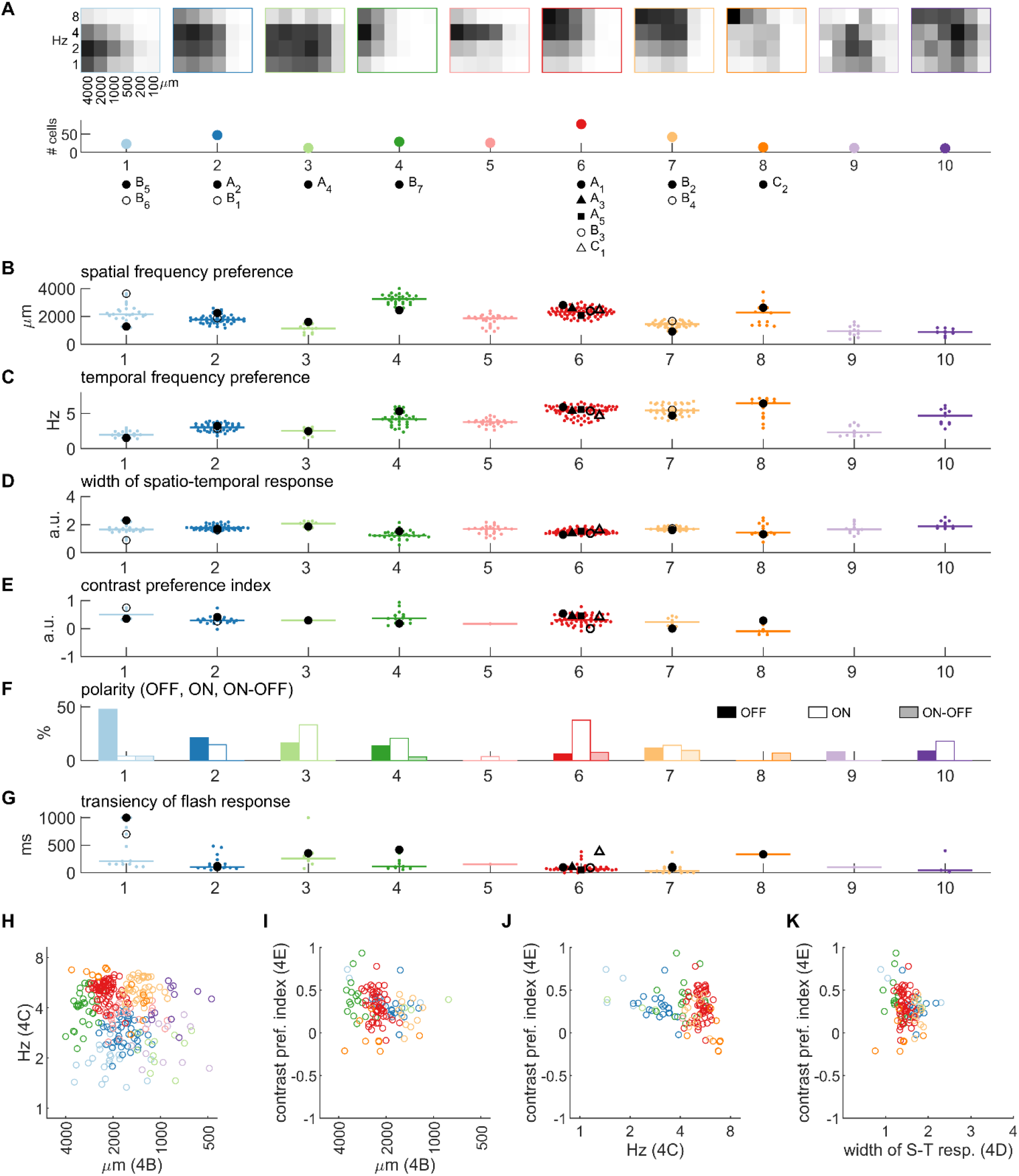
Clusters of human retinal ganglion cells. (A) Heatmaps of spatio-temporal response amplitudes (N = 293) were clustered into 10 groups. For each cluster, the average heatmap is shown as well as the number of cells in that cluster. (B) Preferred spatial frequency based on a Gaussian fitted to the spatio-temporal heatmap. Median and example cells from Figure 2 are shown. (C) Preferred temporal frequency, calculated as for B. (D) Broadness of spatio-temporal response measured as the mean of the two axes of the Gaussian fit. (E) Contrast preference index calculated as the response difference for high vs. low contrast. Responses only to high contrast obtain a value of 1, responses only to low contrast −1. (F) Percentage of OFF, ON, and ON-OFF cells in each cluster. (G) Transciency of flash responses as time from peak to return to background + 2 STD. (H) Preferred spatial and temporal frequency for all cells with responses to drifting-grating. Coloured dots indicate mean per cluster. (I) Preferred spatial frequency and contrast preference index for all cells responding to both drifting-grating and chirp. (J). Preferred temporal frequency and contrast preference index. (K) Width of spatio-temporal response heatmap and contrast preference index.

We found three clusters with responses to a broader range of spatial frequencies and lower temporal frequencies (cluster 1-3, n = 23, 47, 12 cells). Two clusters contain cells with a preference for 4 Hz stimuli, one with cells responding only to wide gratings (cluster 4, n = 29), another one with responses to a broad range of spatial frequencies (cluster 5, n = 26). Clusters 6-8 contain cells that respond best to high temporal frequencies with different spatial preferences (n = 77, 42, 14). Finally, cells in clusters 9 and 10 respond to medium temporal and spatial frequencies (n = 12, 11). While the cells within these clusters showed consistent spatial-temporal tuning to the drifting-grating stimulus, we expect that at least some of these clusters will contain more than one cell type. For instance, cluster 6 contains ON cells, OFF cells, and ON-OFF cells (Fig. 4A). We did not obtain enough responses to other stimuli to include those in the clustering process.

We measured several response parameters to characterize each cluster (Figure 4B to 4G, Table 2), which are summarized and color-coded in Figure 4H to 4K. To more accurately measure the frequency preferences of human ganglion cells, we fit a Gaussian to the response heatmaps (see Methods). Figure 4B shows the location of the peak of this Gaussian along the spatial axis. The temporal preference for each cell per cluster is shown in Figure 4C and the specificity of this spatio-temporal response pattern, i.e. the width of the Gaussian, in Figure 4D. Example cells from Figure 2 are indicated with different markers as listed in Figure 4A.

**Table 2:** Response behavior of clustered human retinal ganglion cells. Mean and standard deviation of the parameters depicted in Figure 4B-G. Spatial and temporal frequency preferences were defined as the peak of a Gaussian fitted to the heatmap of responses to 24 drifting-gratings. The width of this Gaussian is indicated in the column ‘Width S-T’. A contrast preference index was calculated from the responses to the chirp stimulus. An index of 1 indicates responses exclusively to high contrast, an index of −1 exclusive responses to low contrast. Transiency was calculated based on responses to a full-field step. It was defined as the time from the peak response to an activity level ≤ background activity (mean + 2 standard deviations in the 1.5 s before stimulus onset). Contrast preference index and transiency were calculated based on other stimuli than the drifting-gratings used for clustering; the number of cells responding to those stimuli is indicated for each cluster.

| Cluster | N | Spatial frequency preference [μm] (4B) | Temporal frequency preference [Hz] (4C) | Width S-T [a.u.] (4D) | Contrast preference index [a.u.] (4E) | Transiency [ms] (4G) |
| --- | --- | --- | --- | --- | --- | --- |
| 1 | 23 | 2269 ± 597 | 2.0 ± 0.4 | 1.7 ± 0.3 | 0.51 ± 0.21<br>N = 4 | 406 ± 350<br>N = 13 |
| 2 | 47 | 1798 ± 306 | 3.0 ± 0.5 | 1.8 ± 0.2 | 0.30 ± 0.13<br>N = 20 | 171 ± 132<br>N = 17 |
| 3 | 12 | 1130 ± 304 | 2.4 ± 0.5 | 2.0 ± 0.2 | 0.32 ± 0.06<br>N = 3 | 352 ± 333<br>N = 6 |
| 4 | 29 | 3219 ± 358 | 4.2 ± 1.0 | 1.3 ± 0.3 | 0.41 ± 0.22<br>N = 16 | 174 ± 119<br>N = 11 |
| 5 | 26 | 1738 ± 394 | 3.7 ± 0.5 | 1.6 ± 0.3 | 0.17<br>N = 1 | 154<br>N = 1 |
| 6 | 77 | 2317 ± 314 | 5.3 ± 0.8 | 1.5 ± 0.2 | 0.32 ± 0.17<br>N = 54 | 86 ± 73<br>N = 40 |
| 7 | 42 | 1434 ± 216 | 5.5 ± 0.8 | 1.7 ± 0.1 | 0.22 ± 0.16<br>N = 10 | 64 ± 93<br>N = 15 |
| 8 | 14 | 2176 ± 768 | 5.8 ± 1.5 | 1.6 ± 0.5 | -0.04 ± 0.17<br>N = 8 | 335<br>N = 1 |
| 9 | 12 | 965 ± 375 | 2.6 ± 0.8 | 1.7 ± 0.3 | no data | 101<br>N = 1 |
| 10 | 11 | 890 ± 221 | 4.5 ± 1.1 | 2.0 ± 0.3 | no data | 154 ± 214<br>N = 3 |

For every cell that was exposed and responded to a chirp stimulus, we calculated a contrast preference index which is 1 if the cell responded only to high contrast and −1 if it responded only to low contrast (Fig. 4E). The polarity of all cells that responded to full-field flashes is depicted in Figure 4F and the transience of this response, expressed as the time after peak until the response reaches again background + 2 standard deviation firing rates, in Figure 4G.

The most common ganglion cells in the primate retina are midget and parasol cells [9–11]. Midget cells respond to a broad range of spatial frequencies, show sustained responses and low contrast gain [9,39]. In our data set, cluster 1 and 3 could include midget cells. Example cells B_5_ and A_4_ in these two clusters have sustained responses (355 and 1000 ms), a small contrast preference index (0.29 and 0.35) and a rather broad spatio-temporal response profile (2.3 and 1.9 a.u.). Clearly, cluster 1 also contains cells of other types with narrower frequency preferences and stronger responses to high contrast such as example cell B_6_ (contrast preference index = 0.74) as well as cells with transient responses.

Parasol cells, on the other hand, have transient responses, respond well to many spatial frequencies, but especially to wide stimuli, show strong responses to high temporal frequencies, and exhibit a high contrast gain [9,40,41]. Similar stimulus preferences can be seen in our cluster 6. Especially example cells A_1_ and A_3_ behave like classical ON parasol cells (transiency: 100 and 103 ms, contrast preference index: 0.54 and 0.45, spatial frequency preference: 2800 and 2570 μm, temporal frequency preference: 5.9 and 5.4 Hz), and B_3_ could be an OFF parasol cell (transiency: 92 ms, spatial frequency preference: 2400 μm, temporal frequency preference: 5.4 Hz). In addition to putative parasol cells, cluster 6 also contains fast ON-OFF cells such as example C_1_.

Petrusca and colleagues described another ganglion cell type in the primate retina, the upsilon cell, with preferences for big stimuli [24]. The fast cells in cluster 4 (example cell B_7_; transiency: 418 ms, contrast preference index: 0.18, spatial frequency preference: 2440 μm, temporal frequency preference: 5.3 Hz) show a similar behavior as upsilon cells with a preference for wide gratings and >4 Hz stimulation, which is the frequency Petrusca and colleagues found to be optimal to stimulate upsilon cells. Taken together, these observations suggest that we have measured typical responses of the most common primate retinal ganglion cell types.

### Ex-vivo human retinas show physiological properties of healthy tissue

One potential problem when working with human retinas is the unclear health status of the donor tissue. We obtained retinas from donors between 42 and 89 years of age and with different medical histories (Table 1). In addition to the variability introduced by the donors, several circumstances can harm the tissue and prevent light responses: Depending on the surgery procedure, the retina within the ligated eye bulb might have been exposed to longer periods without oxygen and nutrients (ischemia). Furthermore, because of the growing tumor, the retina might have been detached from the pigment epithelium prior to the surgery, which is particularly harmful to photoreceptors. In this study, we thus excluded all retinas exposed to ≥18 min of ischemia (control experiments with ischemic pig eyes have shown a strong decrease in light responses for longer ischemia times; see also [44]). Further, we recorded only from retinal pieces in the opposite hemisphere containing the tumor, and we included in the analysis only retinal pieces from which we could record light responses from at least 10 cells.

We performed several tests to assess the health status of the donor tissue. One hallmark of degenerating retina is tissue-wide oscillatory activity. Such oscillations have been observed in mouse models for retinitis pigmentosa [45] and have a frequency of approximately 9 Hz. In these retinas, each ganglion cell shows oscillatory activity which is synchronized across the whole tissue. We did not observe such oscillations in any of the recorded human retinas. Moreover, light-responsive cells (displayed as green circles overlaid over the MEA electrode grid in Fig. 5A) were distributed across the retinal piece, indicating good recording conditions.

**Figure 5:**
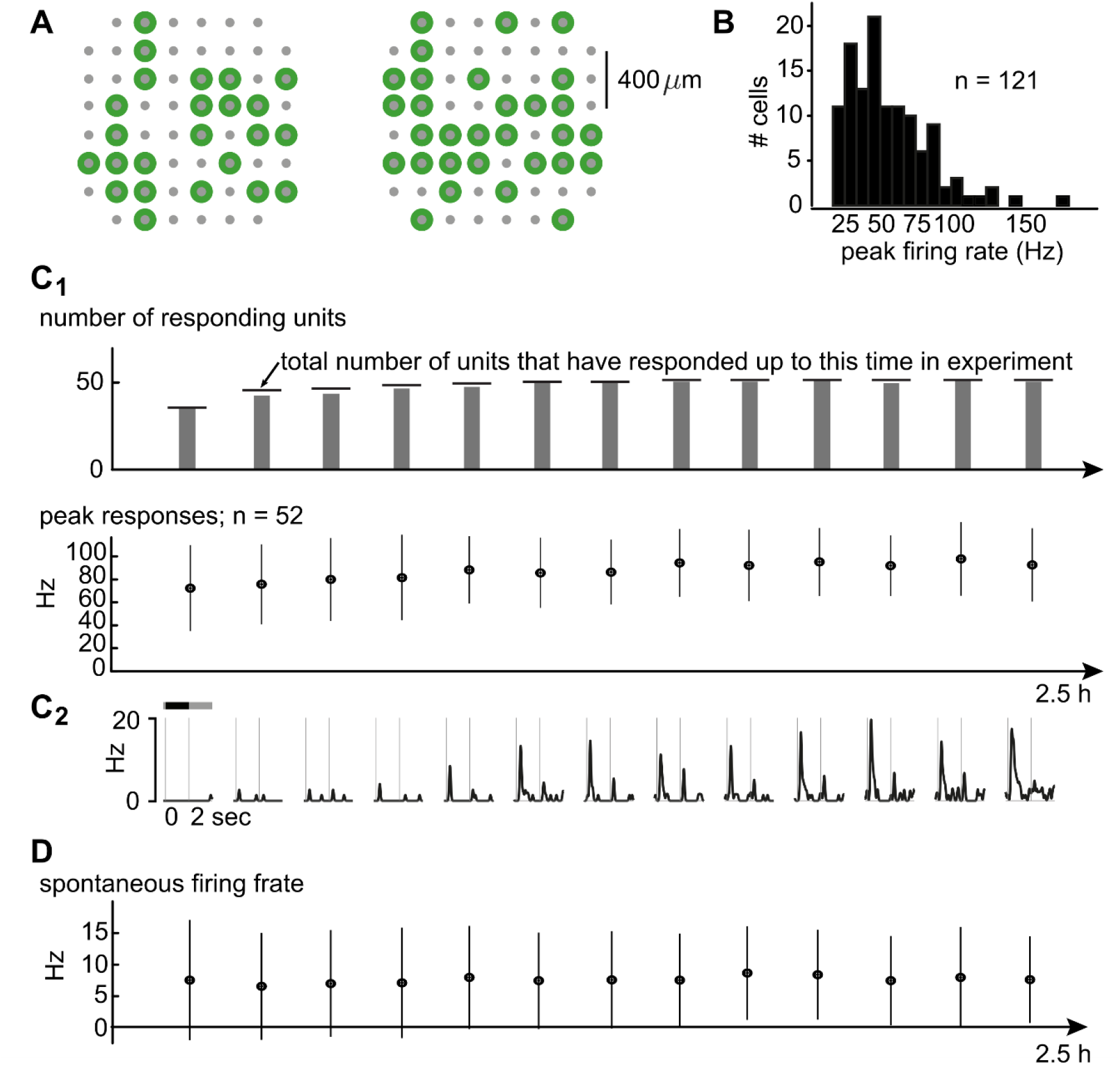
Donated human retinas are healthy. (A) Multi-electrode array layout (gray) and electrodes with sortable, light-responsive cells (green) of two example experiments. Responding cells are distributed across the recorded retinal pieces. (B) Distribution of peak firing rates in response to full-field contrast steps. (C_1_) Top: Number of cells responding at any given time point (gray bars) was similar to the total number that had responses until that time point (horizontal lines). Bottom: Mean ± standard deviation of peak firing rate for the responding cells. Data from a subset of experiments that lasted for 2.5h (N = 4 experiments, N= 52 cells). (C_2_) Example firing rate traces for one cell. (D) Spontaneous background firing rates (mean ± standard deviation) of the same cells and time points as in C_1_.

Overall response strength is another indication of tissue health. We compared the response strength of the recorded cells in the human retina with published primate data, and computed the firing rate in the same way as previous publications on macaque retina [46]. We then extracted the peak firing rate for each cell to the full-field contrast steps. Figure 5B shows the distribution of peak firing rates: many cells produced maximal responses of 20-90 Hz, but peaks could reach up to 180 Hz. Under comparable conditions (binary full-field noise), Uzzell & Chichilnisky report example cells with response peaks between <80 and 300 Hz [46]. The amplitude of the human response peaks reported here is hence in the same range as found in macaque retina.

Peak firing rates were not only comparable to published monkey data but were also stable throughout the experiments. Four retinal pieces were recorded for 2.5 hours and we computed peak responses averaged across blocks of 5 full-field contrast steps across the whole experiments. While some cells did not respond to this stimulus in the very beginning of the experiment, they responded consistently once they started and their peak firing rates were stable (Fig. 5C_1_). The responses at each time point (averaged across blocks of 5 steps) for an example cell are shown in Figure 5C_2_. Similarly, spontaneous background spiking activity was stable across the full recording (Fig. 5D). Taken together, the recorded human retinas did not show typical signs of deteriorating or degenerated tissue and exhibited stable spontaneous and evoked activity.

## Discussion

In this study, we describe light response properties of human retinal ganglion cells and find that these properties are very diverse. We found cells that responded only to positive contrast steps (ON cells), cells that responded only to negative contrast (OFF cells), and cells that encoded both positive and negative contrast (ON-OFF cells). The recorded human ganglion cells preferred different spatio-temporal stimulus frequencies and had distinct response properties when presented with local stimuli or contrast ramps. This diversity is consistent with the variety of cell types predicted by morphological classification in primate retina [11,21]. Our extensive dataset of 324 light-responsive ganglion cells provides an overview of the visual features routed by the human retina to the brain and suggests a similar richness of information processing in the primate retina as found in other mammals where recent studies estimate over 40 distinct retinal information streams [2,3].

### Cell classes in the data set and correspondence with primate literature

Based on spatio-temporal response properties, we identified 10 clusters of retinal ganglion cells. We found response properties typical for midget ganglion cells in clusters 1 and 3 and parasol-like responses in cluster 6. In addition, we found cells in cluster 4 with similar response properties as the previously described upsilon cell [24], a cell type that has been suggested to be an analogue to the cat’s non-linear, fast, and transient Y-cell. Together, these data suggest that it is possible to record previously characterized response properties of non-human retinal ganglion cells also in *ex-vivo* human tissue (see also [18]).

We found midget-like responses in two different clusters and several of the clusters span a broad range for some of the measured response parameters, which indicates that they contain more than one cell type. Varying eccentricity of our recordings might be a reason for the same cell type being assigned to more than one cluster. Especially the spatial frequency preferences of cells will tend to increase with eccentricity due to their increasing size and decreasing cell density. Hence, clustering based on spatio-temporal properties might have assigned midget cells with different spatial preferences to different clusters. On the other hand, clusters may contain more than one cell type because those types may differ in properties other than spatio-temporal preferences. For instance, ON, OFF and ON-OFF cells can be found in most of the ten clusters – a property which cannot be distinguished based on response peaks to drifting-gratings. To further separate cell types with similar spatio-temporal properties from each other, future studies could use denser recordings that allow to test for mosaic formation or other visual stimuli that drive a large population of human ganglion cells.

### Previously unreported response properties in primate retina

Studies of primate retina often aim to characterize in detail a selected type of ganglion cell. In the present study, we considered all cells with light responses, and did not select for specific response features. This approach led to the characterization of response properties that have not been reported previously. The remaining five clusters (5, 7, 8, 9, 10) that do not correspond to previously characterized ganglion cell types contain cells that do not respond well to full-field stimuli (only few cells respond to flashes and/or chirp). Some cells in these clusters, including the example cells B_2_, B_4_ (cluster 7) and C_2_ (cluster 8), responded to a moving bar. The different clusters have different spatio-temporal preferences, from tuning to small, slow stimuli (cluster 9) to a preference for fast and wider stimuli (cluster 8). The lack of responses to full-field stimuli and the specific spatio-temporal preferences suggest that some of these cells are responsible for detection of local stimuli of different size and speed. Further experiments will be necessary to assign those clusters to anatomically or molecularly identified cell types in the primate retina.

### Possibility of overestimation of diversity

There are three main aspects that might have led to an over-estimation of the diversity in our data set. First, most of our stimuli were full-field, and thus responses can reflect center-surround interactions. These interactions can be very diverse across different cell types, such that full field stimuli might help to distinguish cell responses that may otherwise be very similar during local stimulation. The specific surround circuitry can depend strongly on the exact stimulus conditions including stimulus size [47] and absolute light level [47–49]. This dependency can contribute to an overestimation of the diversity of cell types across different recordings. In particular, the diverse history of the donor tissue (age, health and genetic background of the donor, see the third point below), may have consequences for the surround contribution, such that full-field stimulation may exaggerate differing responses in cells of the same type across different recordings. However, under our controlled stimulus conditions, it is not very likely that our large stimuli caused much artificial response variability. What is more, we might even have under-estimated the diversity in the responses as we probably have not recorded from cells that only respond to local stimuli.

Second, variety in the eccentricity of the retinal pieces may have introduced additional diversity in the response properties. This being said, non-human primate studies investigating specific ganglion cell types tend to focus on a large proportion of the retina (e.g. 25-70 degrees in [26], 30-60 degrees in [16]), but did not report any significant differences in response properties across eccentricities.

Third, we cannot exclude that the donor’s health status could have altered the responses of the ganglion cells. Still, on average, the human ganglion cells recorded here showed a similar response behavior to drifting-grating stimuli of different temporal and spatial frequencies as previously published non-human primate ganglion cells. The most abundant cell types in the primate retina are midget and parasol cells. Our dataset contains cells that responded similarly stimuli as previously described midget and parasol cells in other primates. Furthermore, the spatial threshold for the whole population of recorded cells (1.55 cyc/°) is comparable to psychophysically determined spatial resolution thresholds of human subjects measured at comparable eccentricity. This, together with the absence of oscillations and the fact that we observed responding ganglion cells distributed across many recording electrodes, suggests that we were able to record physiologically relevant response properties in these donor human retinas.

### Absence of direction-selective cells

One of the best studied ganglion cells in non-primate mammalian retina are the direction-selective ganglion cells. It is unclear whether cells responding to a specific direction of movement exist in the primate retina. So far, no physiological recordings of direction-selective cells have been published (but see [50]) and we could not detect such direction-selective behavior in our data set either (data not shown). Morphological studies identified potential candidates for primate direction-selective neurons [23,51,52]. These cells have a large dendritic field and hence they are much fewer in number than the smaller midget or parasol cells. Consequently, the chances to record from such large cells in unbiased MEA experiments is small. Furthermore, as shown in the present study as well as in previous measurements [53], primate ganglion cells respond to higher temporal frequencies than for instance mouse ganglion cells [54]. It is therefore possible that our and other studies missed direction-selective cells in primate retinas due to suboptimal stimulation paradigms. This should be taken into consideration for future studies.

### Future studies on the output of the human retina

In this study we provide the first non-selective description of the retinal output in humans. We showed that our data is consistent with measurements in other primates and that the diversity in the human retinal output is larger than suggested by previous physiological studies that focused on only a few primate retinal cell types. To further investigate these unstudied ganglion cells and to achieve classification into individual cell types, one could make use of high-density MEAs. These MEAs allow recording from almost every cell in a given patch [32], and local stimulation would be possible to test for parameters such as center-surround mechanisms, local edge detection or approach sensitivity. It has been shown that each ganglion cell type tiles the retina with little overlap in order to encode every visual feature at each point in the visual field [55]. Such mosaic formation can as well be revealed with high-density MEA recordings [15,16,24] and can then be used for cell type identification.

### Impact on bio-medical research

The goal of bio-medical research is to better understand human physiology and to find treatments in the case of disease. Knowledge about the detailed functioning of the human retina would be desirable also in the context of retinal diseases. Such diseases, in particular blindness, have a big impact on individuals and the society. In recent years, research has yielded some promising approaches to potentially healing blindness (e.g. electrical retinal implants [56,57], optogenetics [58,59], stem cell therapy [60]) with a common ultimate goal: to come as close as possible to full vision capabilities by interfering appropriately with the retina of the patient. Especially optogenes (light sensitive ion channels/pumps) are a promising tool to render degenerated photoreceptors, bipolar cells, or ganglion cells light sensitive [58,59,61–63]. Currently, these treatment options are mostly developed and tested in animal models. We see a big advantage of supplementing this research with human retina studies. First, increased knowledge about signal processing within the human retina may support further and faster progress in that field. Second, cell type specificity of viral vectors and the correct expression of the genetic construct containing the optogenes could be developed using *ex-vivo* or *post-mortem* human retina. Moreover, by subsequent comparison of the optogene-driven light responses with the natural responses presented in this and future studies, one could evaluate the efficacy of the treatment. Finally, (side-)effects of drugs such as neuroprotectiva (substances to conserve as much as possible of leftover visual capabilities) could be tested directly on human retina instead of using porcine, bovine or other animal models. First studies have started using human tissue to characterize the genetic profile of human retinal cells and to test first treatment options [64,65]. We hope that the present study may serve as encouragement for more research with *ex-vivo* human retina in the future.

## Methods

Code to recreate several figures and the necessary spike times and processed data can be accessed on GitHub (https://github.com/katjaReinhard/HumRet) and spike times as well as processed data on OSF (https://osf.io/zf9rd/).

### Human retina donations

To characterize information processing in the retina, very fresh tissue is necessary because the photoreceptors rapidly lose light-sensitivity. We obtained such human retina from patients of the University Eye Hospital in Tübingen, who had to undergo enucleation of one eye, usually to remove a tumor. All procedures were approved by the ethics committee of the University of Tübingen (approval number 531/2011) and performed in accordance with the guidelines and regulations provided by the ethics committee. All participants provided informed consent to the use of the removed retina for scientific research purposes. The retina was protected from light during surgery if possible. An ischemia time of at least five minutes during the surgery (clamping of the optic nerve before removing the bulbus) was mandatory to prevent strong bleeding. The bulbus was cut in halves directly after enucleation, and the hemisphere without tumor was put immediately into CO_2_-independent culture medium (Gibco, ThermoFisher Scientific, Massachusetts, USA), kept in darkness at room temperature and transported to our lab. Under dim red light, we removed the vitreous and cut small mid-peripheral retinal pieces (~ 3×3 mm²). Within 23 months we obtained 15 such ex-vivo donations (Table 1). 15 pieces from 10 retinas were used for experiments.

### Experimental design

To maximize the amount of information gained from the rare experiments with fresh human retina, we employed recordings with flat multi-electrode arrays (MEA) that allow for measuring the activity of many neurons in parallel [37]. MEAs include a square or rectangular electrode arrangement that is brought in contact with the ganglion cells, allowing measuring the retinal output in response to light stimulation. Our MEA experiments have been described in detail elsewhere [66]. Briefly, the retinal pieces were placed ganglion cell side-down on a MEA. We used perforated 60-electrode MEAs with 200 μm distance between the electrodes (60pMEA200/30iR-Ti-gr, Multichannel Systems, Reutlingen, Germany). Then, various light stimuli were focused onto the photoreceptors with a Digital Light Processing projector (Sharp PG-F212X-L, Sharp Corporation, Osaka, Japan or Acer K11, Acer, Taipeh, Taiwan), and we recorded the output of the retina (i.e. the action potentials of ganglion cells in response to the stimuli) at 25 kHz with a USB-MEA-system (USB-MEA1060, Multichannel Systems) or an MC-Card based MEA-system (MEA1060, Multichannel Systems). During the experiments, the retina was kept at 25°C and continuously superfused with Ringer solution (in mM: 110 NaCl, 2.5 KCl, 1 CaCl_2_, 1.6 MgCl_2_, 10 D-Glucose, and 22 NaHCO_3_; ~270 mosm) or modified Ringer solution (in mM: 115 NaCl, 2.5 KCl, 2 CaCl_2_, 1 MgCl_2_, 15 D-Glucose, 1.3 NaH_2_PO_4_*H_2_O, 0.5 L-Glutamine, and 25 NaHCO_3_; ~285 mosm), both equilibrated with carbogen (95% O_2_, 5% CO_2_). All experiments were conducted with the retinal pigment epithelium removed.

### Light stimulation

The stimulation intensity provided by our projectors spanned 3 log units of brightness between a black (‘0’) and white (‘255’) stimulus. The projector output was linearized, so that the grey (‘128’) background was midway between black and white, and the intensity step between black and grey and between grey and white had equal amplitude. Recordings were performed at photopic intensity levels (light intensity for day vision) with a mean illuminance of 8·10^4^ rod isomerizations per rod per second. In two retinal pieces, no clear responses could be detected at this light level, and we used data obtained at a mean illuminance of 8·10^5^ rod isomerizations per rod per second for analysis. Note that recordings at photopic light levels do not necessarily imply that the observed light responses were driven by cones alone, rods may have contributed as well [67]. A broad set of light stimuli was used; each stimulus was repeated several times during recording sessions of two to six hours. We calculated various parameters from the ganglion cells’ responses (see below). To convert stimulus sizes on the retina (in μm) to the equivalent visual angles (in degree), we used the conversion factor 266 μm/° [68]. We discuss in this article six response parameters extracted from responses to the following six stimuli (see also Fig. 1):

#### Sinusoidal drifting-gratings

Drifting sinusoidal grating stimuli with 24 different combinations of spatial periods and temporal frequencies (1, 2, 4, 8 Hz; 100, 200, 500, 1000, 2000, 4000 μm spatial period on the retina) were used for spatio-temporal analysis (Fig. 1A). The gratings were shown at full contrast (‘0’ to ‘255’) and moved in one direction for 12 seconds.

#### Temporal and contrast chirp

Temporal tuning was also tested with a spatially homogeneous chirp stimulus [2], i.e. full-field frequency-modulated intensity change between black (‘0) and white (‘255’), according to: *intensity* = 128 + 128 * sin(*π*(*t*^2^ + *t*/10)), with t given in seconds. The temporal frequency increased from 0.5 to 8 Hz over a time course of approximately 8 seconds (Fig. 1B top). In addition, a contrast ramp increasing from none to full contrast within 8 seconds was used to test for contrast sensitivity (Fig. 1B bottom).

#### Single bars at various velocities

We used single bars moving with different speed to test for speed preferences. A bar with 1000 μm extension in the movement direction (either black or white) and covering the complete screen in the other direction moved in front of a gray background in one direction (same direction as grating stimulus) with different speeds (1, 2, 4, 8, 16 mm/s) with a gap of 3 seconds before the next higher speed (Fig. 1C).

#### Full-field contrast steps

Full-field contrast steps were applied for measurements of response polarity and latency (Fig. 1D). A single stimulus consisted of four transitions (grey → black → grey → white → grey) spanning the full projector intensity of 3 log units of brightness (contrast for each step: ± 1 Weber contrast). Each contrast step lasted for 2 seconds.

#### Direction-selectivity

We used a single bar (black or white) moving in 8 directions to test for direction-selectivity. The bar of 1000 μm width was moved with 1 mm/s across the retina.

### Spike extraction

Spike sorting (assignment of single action potentials to individual cells) was performed with an in-house Matlab (MathWorks, Massachusetts, USA) routine written by Alexandra Kling. Different features of the action potential waveforms, such as amplitude, width, or principal components, were calculated and projected onto 2-dimensional space to separate action potentials of different cells from each other and from noise. In addition, the spike refractory time of all spikes of a sorted cell had to be >1.5 ms. After spike sorting, we determined light-responding cells by visual inspection of the activity to all stimuli. To calculate the firing rate, the spike train was convolved with a Gaussian and plotted against time. The sigma of the Gaussian varied for different analysis purposes; the value applied in each case is given in the description below. For the firing rates of the example cells in Figure 2, σ = 40 ms was used. Only cells for which spikes could be sorted confidently were used for analysis (for consistency, the same person performed spike sorting for all experiments and applied the identical quality judgement system). We applied cross-correlation analysis to detect recordings from the same cell on different electrodes (e.g. from cell body and axon). In this case, only one of the recorded units was used for the analysis.

### Response parameter calculation

#### Spatio-temporal tuning

Drifting sinusoidal grating stimuli were used for spatio-temporal analysis. First, cells responding to at least one of the drifting gratings were identified manually. For each cell and stimulus repetition we represented the cell’s activity with a binary vector indicating the presence (1) or absence (0) of a spike (time bins: 1 ms). For each drifting grating stimulus, we then calculated the mean of these binary spike rates and computed its Fourier transform (FT). The FT peak at the stimulus frequency was then taken as the cell’s response strength. The Fourier transform was considered to have a peak (i.e., the cell was considered to respond to the stimulus) if there was no higher peak at any other frequencies (excluding multiples of the stimulus frequency).

#### Temporal tuning

Temporal tuning was tested with a chirp stimulus, i.e. frequency-modulated sinusoidal full-field change of intensity. We calculated the FT of both, the stimulus and the response (mean binary spike train, frequency resolution of 0.125 Hz). Response strength along the stimulation frequencies was defined as *norm*(*FT*_*response*_)/*FT*_*stimulus*_. Fluctuations were smoothed; these appeared especially at low temporal frequencies due to the timing of ON- and OFF-responses. Smoothing was achieved by averaging of the response strength with a moving average across a 3-datapoint-window (0.375 Hz) in steps of 1 data-point (0.125 Hz). Population data is presented in 2.6 Hz bins across all responding cells (Fig. 2C). As a second method, temporal tuning was also calculated from the FT amplitudes obtained from the responses to drifting-grating stimuli.

#### Median speed preference

A black or white bar was moved across the retina in one direction (same direction as drifting granting) with various speeds. The cumulative sum of peak responses for each speed (firing rate calculated with σ = 40 ms) was computed. The speed value for which 50% of the cumulative sum was reached was taken as the cells’ median speed preference. For each cell that responded to both, white and black bars, the higher preferred speed was taken for the population plot in Figure 2D.

#### Polarity

Polarity was defined based on the responses to full-field contrast steps. Cells with responses only for positive contrast steps were considered as ON-cells; OFF-cells had only detectable responses to negative contrasts, and ON-OFF-cells responded to both types of contrast steps. Firing rates were calculated by convolving the spike rates with a Gaussian (σ = 40 ms). The cell was considered to show a response if the peak firing rate was bigger than mean spontaneous activity + 2 standard deviations (measured before the first step in contrast).

#### Transiency

Response transiency was defined as the time between the peak response to full-field contrast steps and returning of the spiking activity to baseline firing rate + 2 standard deviations. Shorter times correspond to more transient responses.

#### Clustering of spatio-temporal responses

The 24 peak responses were normalized for each cell so that the response to the optimal drifting-grating was 1. These normalized responses were clustered for the 293 responding cells using k-means in MATLAB (squared Euclidean distance, 1000 repetitions). The best number of clusters was identified using the Calinski-Harabasz [69] and Davies-Bouldin indices [70] (‘evalclusters’ in MATLAB).

#### Gaussian fit to spatio-temporal heat-maps

We fit a Gaussian to the spatio-temporal response heatmaps to calculate frequency preferences and specificity of the response. The underlying code can be found on GitHub (https://github.com/katjaReinhard/HumRet). The center point and hence the preferred overall spatial and temporal frequencies were defined as the dot product of the heatmap with a mesh of spatial and temporal frequencies, respectively. The dot product of the squared distance of each heatmap (SF for spatial, TF for temporal frequency) point with the Gaussian center (Csf, Ctf) and the heatmap itself (peaks) was then calculated (*a* = (*SF* − *Csf*)^2^ · *peaks*, *b* = ((*TF* − *Ctf*) * (*SP* − *Csf*)) · *peaks*, *c* = (*TF* − *Ctf*)^2^ · *peaks*). The squared Eigenvalues of [a,b;b,c] were considered the major axes of the Gaussian and their average was taken as the spatio-temporal response width. A bigger width indicates a less specific preference.

#### Contrast preference index

To compute the contrast preference index, the first third (low contrast) and last third (high contrast) of the background-subtracted response to the contrast part of the chirp stimulus was taken. The index was calculated as *CPI* = (*high* − *low*)/(*high* + *low*) with ‘high’ indicating the sum of the absolute firing rate during high contrast and ‘low’ the sum of the absolute firing rate during low contrast.

## Acknowledgements

We thank Prof. Karl Ulrich Bartz-Schmidt and the surgery team of the University Clinics Tübingen for the support during the human retina donation process. We further thank Dr. Hartwig Seitter (University of Innsbruck, Austria), Dr. Boris Benkner, Dr. Marion Mutter, and Natalia Świętek for technical support; Dr. Alexandra Kling (Stanford University, California, USA) for the spike-sorting code; and Dr. Karl Farrow and Dr. Vincent Bonin (both NERF, Leuven, Belgium) for helpful comments on the manuscript. This research was supported by funds of the Deutsche Forschungsgemeinschaft (DFG) to the Werner Reichardt Centre for Integrative Neuroscience (DFG EXC 307), by the Bundesministerium für Bildung and Forschung (BMBF) to the Bernstein Center for Computational Neuroscience (FKZ 01GQ1002), a Pro-Retina Stipend to KR, and the Lush Prize 2015 for young investigators to KR.

## References

1. Lettvin JY, Maturana HR, Maturana HR, Mcculloch WS, Pitts WH. What the Frog’s Eye Tells the Frog’s Brain. Proc IRE. 1959;47: 1940–1951. doi:10.1109/JRPROC.1959.287207

2. Baden T, Berens P, Franke K, Roman-Roson M, Bethge M, Euler. The functional diversity of mouse retinal ganglion cells. Nature. 2016;529: 1–21. doi:10.1038/nature16468

3. Sanes JR, Masland RH. The types of retinal ganglion cells: current status and implications for neuronal classification. Annu Rev Neurosci. 2015;38: 221–46. doi:10.1146/annurev-neuro-071714-034120

4. Farrow K, Masland RH. Physiological clustering of visual channels in the mouse retina. J Neurophysiol. 2011/01/29. 2011;105: 1516–1530. doi:10.1152/jn.00331.2010

5. Devries SH, Baylor DA. Mosaic arrangement of ganglion cell receptive fields in rabbit retina. J Neurophysiol. 1997/10/27. 1997;78: 2048–2060. Available: http://www.ncbi.nlm.nih.gov/pubmed/9325372

6. Wong RC, Cloherty SL, Ibbotson MR, O’Brien BJ. Intrinsic physiological properties of rat retinal ganglion cells with a comparative analysis. J Neurophysiol. 2012/07/13. 2012;108: 2008–2023. doi:10.1152/jn.01091.2011

7. Marre O, Amodei D, Deshmukh N, Sadeghi K, Soo F, Holy TE, et al. Mapping a complete neural population in the retina. J Neurosci. 2012/10/27. 2012;32: 14859–14873. doi:10.1523/JNEUROSCI.0723-12.2012

8. Polyak SL. The Retina. Illinois: The University of Chicago; 1941.

9. Kaplan E, Shapley RM. The primate retina contains two types of ganglion cells, with high and low contrast sensitivity. Proc Natl Acad Sci U S A. 1986;83: 2755–7. Available: http://www.ncbi.nlm.nih.gov/pubmed/3458235

10. Rodieck RW, Binmoeller KF, Dineen J. Parasol and midget ganglion cells of the human retina. J Comp Neurol. 1985;233: 115–132. doi:10.1002/cne.902330107

11. Dacey DM. Origins of perception: retinal ganglion cell diversity and the creation of parallel visual pathways. The Cognitive Neurosicences. 2004. pp. 281–301.

12. Chichilnisky EJ, Kalmar RS. Functional asymmetries in ON and OFF ganglion cells of primate retina. J Neurosci. 2002;22: 2737–47. doi:20026215

13. Dacey DM, Brace S. A coupled network for parasol but not midget ganglion cells in the primate retina. Vis Neurosci. 1992;9: 279–290. doi:10.1017/S0952523800010695

14. Dacey DM, Lee BB. The “blue-on” opponent pathway in primate retina originates from a distinct bistratified ganglion cell type. Nature. 1994;367: 731–735. doi:10.1038/367731a0

15. Gauthier JL, Field GD, Sher A, Shlens J, Greschner M, Litke AM, et al. Uniform Signal Redundancy of Parasol and Midget Ganglion Cells in Primate Retina. J Neurosci. 2009;29: 4675–4680. doi:10.1523/JNEUROSCI.5294-08.2009

16. Greschner M, Shlens J, Bakolitsa C, Field GD, Gauthier JL, Jepson LH, et al. Correlated firing among major ganglion cell types in primate retina. J Physiol. 2011;589: 75–86. doi:10.1113/jphysiol.2010.193888

17. Shlens J, Field GD, Gauthier JL, Greschner M, Sher A, Litke AM, et al. The Structure of Large-Scale Synchronized Firing in Primate Retina. J Neurosci. 2009;29: 5022–5031. doi:10.1523/JNEUROSCI.5187-08.2009

18. Soto F, Hsiang JC, Rajagopal R, Piggott K, Harocopos GJ, Couch SM, et al. Efficient Coding by Midget and Parasol Ganglion Cells in the Human Retina. Neuron. 2020 [cited 22 Jul 2020]. doi:10.1016/j.neuron.2020.05.030

19. Kling A, Gogliettino AR, Shah NP, Wu EG, Brackbill N, Sher A, et al. Functional Organization of Midget and Parasol Ganglion Cells in the Human Retina. bioRxiv. 2020; 2020.08.07.240762. doi:10.1101/2020.08.07.240762

20. Doi E, Gauthier JL, Field GD, Shlens J, Sher A, Greschner M, et al. Efficient Coding of Spatial Information in the Primate Retina. J Neurosci. 2012;32: 16256–16264. doi:10.1523/JNEUROSCI.4036-12.2012

21. Field GD, Chichilnisky EJ. Information Processing in the Primate Retina: Circuitry and Coding. Annu Rev Neurosci. 2007;30: 1–30. doi:10.1146/annurev.neuro.30.051606.094252

22. Peterson BB, Dacey DM. Morphology of human retinal ganglion cells with intraretinal axon collaterals. Vis Neurosci. 1998;15: 377–87. Available: http://www.ncbi.nlm.nih.gov/pubmed/9605537

23. Yamada ES, Bordt AS, Marshak DW. Wide-field ganglion cells in macaque retinas. Vis Neurosci. 2005;22: 383–393. doi:10.1017/S095252380522401X

24. Petrusca D, Grivich MI, Sher A, Field GD, Gauthier JL, Greschner M, et al. Identification and characterization of a Y-like primate retinal ganglion cell type. J Neurosci. 2007;27: 11019–27. doi:10.1523/JNEUROSCI.2836-07.2007

25. Chichilnisky EJ, Baylor DA. Receptive-field microstructure of blue-yellow ganglion cells in primate retina. Nat Neurosci. 1999;2: 889–893. doi:10.1038/13189

26. Field GD, Sher A, Gauthier JL, Greschner M, Shlens J, Litke AM, et al. Spatial Properties and Functional Organization of Small Bistratified Ganglion Cells in Primate Retina. J Neurosci. 2007;27: 13261–13272. doi:10.1523/JNEUROSCI.3437-07.2007

27. Dacey DM, Liao H-W, Peterson BB, Robinson FR, Smith VC, Pokorny J, et al. Melanopsin-expressing ganglion cells in primate retina signal colour and irradiance and project to the LGN. Nature. 2005;433: 749–754. doi:10.1038/nature03387

28. De Monasterio FM, Gouras P. Functional properties of ganglion cells of the rhesus monkey retina. J Physiol. 1975;251: 167–95. Available: http://www.ncbi.nlm.nih.gov/pubmed/810576

29. Schiller PH, Malpeli JG. Properties and tectal projections of monkey retinal ganglion cells. J Neurophysiol. 1977;40: 428–45. Available: http://www.ncbi.nlm.nih.gov/pubmed/403252

30. Hashimoto T, Katai S, Saito Y, Kobayashi F, Goto T. ON and OFF channels in human retinal ganglion cells. J Physiol. 2013;591: 327–337. doi:10.1113/jphysiol.2012.243683

31. Weinstein GW, Hobson RR, Baker FH. Extracellular recordings from human retinal ganglion cells. Science. 1971;171: 1021–2. Available: http://www.ncbi.nlm.nih.gov/pubmed/5542805

32. Fiscella M, Farrow K, Jones IL, Jäckel D, Müller J, Frey U, et al. Recording from defined populations of retinal ganglion cells using a high-density CMOS-integrated microelectrode array with real-time switchable electrode selection. J Neurosci Methods. 2012;211: 103–113. doi:10.1016/j.jneumeth.2012.08.017

33. Frey U, Egert U, Heer F, Hafizovic S, Hierlemann A. Microelectronic system for high-resolution mapping of extracellular electric fields applied to brain slices. Biosens Bioelectron. 2009;24: 2191–2198. doi:10.1016/j.bios.2008.11.028

34. Segev R, Goodhouse J, Puchalla J, Berry MJ. Recording spikes from a large fraction of the ganglion cells in a retinal patch. Nat Neurosci. 2004;7: 1154–61. doi:10.1038/nn1323

35. Zeck GM, Masland RH. Spike train signatures of retinal ganglion cell types. Eur J Neurosci. 2007;26: 367–380. doi:10.1111/j.1460-9568.2007.05670.x

36. Meister M, Lagnado L, Baylor DA. Concerted signaling by retinal ganglion cells. Science. 1995;270: 1207–10. Available: http://www.ncbi.nlm.nih.gov/pubmed/7502047

37. Meister M, Pine J, Baylor DA. Multi-neuronal signals from the retina: acquisition and analysis. J Neurosci Methods. 1994;51: 95–106. Available: http://www.ncbi.nlm.nih.gov/pubmed/8189755

38. Rovamo J, Virsu V, Näsänen R. Cortical magnification factor predicts the photopic contrast sensitivity of peripheral vision. Nature. 1978;271: 54–6. Available: http://www.ncbi.nlm.nih.gov/pubmed/625324

39. Diller L, Packer OS, Verweij J, McMahon MJ, Williams DR, Dacey DM. L and M cone contributions to the midget and parasol ganglion cell receptive fields of macaque monkey retina. J Neurosci. 2004;24: 1079–88. doi:10.1523/JNEUROSCI.3828-03.2004

40. Crook JD, Davenport CM, Peterson BB, Packer OS, Detwiler PB, Dacey DM. Parallel ON and OFF Cone Bipolar Inputs Establish Spatially Coextensive Receptive Field Structure of Blue-Yellow Ganglion Cells in Primate Retina. J Neurosci. 2009;29: 8372–8387. doi:10.1523/JNEUROSCI.1218-09.2009

41. Crook JD, Packer OS, Dacey DM. A synaptic signature for ON- and OFF-center parasol ganglion cells of the primate retina. Vis Neurosci. 2014;31: 57–84. doi:10.1017/S0952523813000461

42. Dacey DM, Crook JD, Packer OS. Distinct synaptic mechanisms create parallel S-ON and S-OFF color opponent pathways in the primate retina. Vis Neurosci. 2014;31: 139–151. doi:10.1017/S0952523813000230

43. Crook JD, Packer OS, Dacey DM. A synaptic signature for ON- and OFF-center parasol ganglion cells of the primate retina. Vis Neurosci. 2014;31: 57–84. doi:10.1017/S0952523813000461

44. Reinhard K, Mutter M, Gustafsson E, Gustafsson L, Vaegler M, Schultheiss M, et al. Hypothermia Promotes Survival of Ischemic Retinal Ganglion Cells. Investig Opthalmology Vis Sci. 2016;57: 658. doi:10.1167/iovs.15-17751

45. Menzler J, Zeck G. Network Oscillations in Rod-Degenerated Mouse Retinas. J Neurosci. 2011;31: 2280–2291. doi:10.1523/JNEUROSCI.4238-10.2011

46. Uzzell VJ, Chichilnisky EJ. Precision of Spike Trains in Primate Retinal Ganglion Cells. J Neurophysiol. 2004;92: 780–789. doi:10.1152/jn.01171.2003

47. Farrow K, Teixeira M, Szikra T, Viney TJ, Balint K, Yonehara K, et al. Ambient illumination toggles a neuronal circuit switch in the retina and visual perception at cone threshold. Neuron. 2013/04/02. 2013;78: 325–338. doi:10.1016/j.neuron.2013.02.014

48. Field GD, Greschner M, Gauthier JL, Rangel C, Shlens J, Sher A, et al. High-sensitivity rod photoreceptor input to the blue-yellow color opponent pathway in macaque retina. Nat Neurosci. 2009;12: 1159–64. doi:10.1038/nn.2353

49. Tikidji-Hamburyan A, Reinhard K, Seitter H, Hovhannisyan A, Procyk CA, Allen AE, et al. Retinal output changes qualitatively with every change in ambient illuminance. Nat Neurosci. 2015;18: 66–74. doi:10.1038/nn.3891

50. Detwiler PB, Crook J, Packer O, Robinson F, Dacey DM. The recursive bistratified ganglion cell type of the macaque monkey retina is ON-OFF direction seletctive. Investigative ophthalmology & visual science. 2019. p. 3884.

51. Moritoh S, Komatsu Y, Yamamori T, Koizumi A. Diversity of Retinal Ganglion Cells Identified by Transient GFP Transfection in Organotypic Tissue Culture of Adult Marmoset Monkey Retina. Chiao C-C, editor. PLoS One. 2013;8: e54667. doi:10.1371/journal.pone.0054667

52. Masri RA, Percival KA, Koizumi A, Martin PR, Grünert U. Survey of retinal ganglion cell morphology in marmoset. J Comp Neurol. 2019;527: 236–258. doi:10.1002/cne.24157

53. Lee BB, Pokorny J, Smith VC, Kremers J. Responses to pulses and sinusoids in macaque ganglion cells. Vision Res. 1994;34: 3081–96. Available: http://www.ncbi.nlm.nih.gov/pubmed/7975341

54. Grubb MS, Thompson ID. Visual Response Properties of Burst and Tonic Firing in the Mouse Dorsal Lateral Geniculate Nucleus. J Neurophysiol. 2005;93: 3224–3247. doi:10.1152/jn.00445.2004

55. Wässle H, Riemann HJ. The mosaic of nerve cells in the mammalian retina. Proc R Soc London Ser B, Biol Sci. 1978;200: 441–61. Available: http://www.ncbi.nlm.nih.gov/pubmed/26058

56. Dorn JD, Ahuja AK, Caspi A, da Cruz L, Dagnelie G, Sahel J-A, et al. The Detection of Motion by Blind Subjects With the Epiretinal 60-Electrode (Argus II) Retinal Prosthesis. JAMA Ophthalmol. 2013;131: 183. doi:10.1001/2013.jamaophthalmol.221

57. Zrenner E, Bartz-Schmidt KU, Benav H, Besch D, Bruckmann A, Gabel V-P, et al. Subretinal electronic chips allow blind patients to read letters and combine them to words. Proc R Soc B Biol Sci. 2011;278: 1489–1497. doi:10.1098/rspb.2010.1747

58. Busskamp V, Roska B. Optogenetic approaches to restoring visual function in retinitis pigmentosa. Curr Opin Neurobiol. 2011;21: 942–946. doi:10.1016/j.conb.2011.06.001

59. Lagali PS, Balya D, Awatramani GB, Münch TA, Kim DS, Busskamp V, et al. Light-activated channels targeted to ON bipolar cells restore visual function in retinal degeneration. Nat Neurosci. 2008;11: 667–675. doi:10.1038/nn.2117

60. Tibbetts MD, Samuel MA, Chang TS, Ho AC. Stem cell therapy for retinal disease. Curr Opin Ophthalmol. 2012;23: 226–34. doi:10.1097/ICU.0b013e328352407d

61. Bi A, Cui J, Ma Y-P, Olshevskaya E, Pu M, Dizhoor AM, et al. Ectopic Expression of a Microbial-Type Rhodopsin Restores Visual Responses in Mice with Photoreceptor Degeneration. Neuron. 2006;50: 23–33. doi:10.1016/j.neuron.2006.02.026

62. Mutter M, Münch TA. Strategies for Expanding the Operational Range of Channelrhodopsin in Optogenetic Vision. Bene F Del, editor. PLoS One. 2013;8: e81278. doi:10.1371/journal.pone.0081278

63. van Wyk M, Pielecka-Fortuna J, Löwel S, Kleinlogel S. Restoring the ON Switch in Blind Retinas: Opto-mGluR6, a Next-Generation, Cell-Tailored Optogenetic Tool. Harris WA, editor. PLOS Biol. 2015;13: e1002143. doi:10.1371/journal.pbio.1002143

64. Nelidova D, Morikawa RK, Cowan CS, Raics Z, Goldblum D, Scholl HPN, et al. Restoring light sensitivity using tunable near-infrared sensors. Science. 2020;368: 1108–1113. doi:10.1126/science.aaz5887

65. Cowan CS, Renner M, Gross-Scherf B, Goldblum D, Munz M, Krol J, et al. Cell types of the human retina and its organoids at single-cell resolution: developmental convergence, transcriptomic identity, and disease map. bioRxiv. 2019; 703348. doi:10.1101/703348

66. Reinhard K, Tikidji-Hamburyan A, Seitter H, Idrees S, Mutter M, Benkner B, et al. Step-By-Step Instructions for Retina Recordings with Perforated Multi Electrode Arrays. Barnes S, editor. PLoS One. 2014;9: e106148. doi:10.1371/journal.pone.0106148

67. Tikidji-Hamburyan A, Reinhard K, Storchi R, Dietter J, Seitter H, Davis KE, et al. Rods progressively escape saturation to drive visual responses in daylight conditions. Nat Commun. 2017;8: 1813. doi:10.1038/s41467-017-01816-6

68. Drasdo N, Fowler CW. Non-linear projection of the retinal image in a wide-angle schematic eye. Br J Ophthalmol. 1974;58: 709–14. Available: http://www.ncbi.nlm.nih.gov/pubmed/4433482

69. Calinksi T, Harabasz J. A dendritic method for cluster analysis. Comm Stat. 1974;3: 1–27.

70. Davies DL, Bouldin DW. A Cluster Separation Measure. IEEE Trans Pattern Anal Mach Intell. 1979;PAMI-1: 224–227. doi:10.1109/TPAMI.1979.4766909

